# Transcranial direct current stimulation over the frontal eye field has no effect on visual search performance

**DOI:** 10.1101/2025.05.27.656327

**Authors:** Charline Peylo, Constanze Albrecht, Ruth Gerlinde Homann, Joshua Richter, Marie Anaïs Dornier, Mona Sophie Ehrat, Finia Luca Loeb, Sophia Manz, Marina Praguer Gaeta, Christina Rieger, Julia Christine Tafelmaier, Paul Sauseng

## Abstract

Top-down attention for the goal-directed (de-)prioritization of information is fundamental for successful everyday-life behavior and poses tremendous problems when negatively impacted by disease. Attention-targeting enhancement and rehabilitation attempts using non-invasive brain stimulation techniques like transcranial direct current stimulation (tDCS) are therefore of major importance. tDCS-driven excitation of the left frontal eye field (FEF; a key region within fronto-parietal attention networks) has recently been suggested to improve attention-guided visual search with stronger effects for lower baseline performers. Here, we report two preregistered tDCS experiments that tested 1) whether the previously observed visual search improvement could be boosted through stimulation over the allegedly more dominant right FEF and 2) whether tDCS-related visual search improvements might depend on search field size. To this end, in experiments one and two, *N*=29 and *N*=31 healthy participants performed a visual search task, in which they searched for an upside-down ‘T’ amongst upright ‘T’s and ‘L’s within small or large search fields, before and during the application of anodal (excitatory) or sham (control) tDCS over the right or left FEF, respectively. In contrast to previous studies, in both experiments (i.e., independent of stimulation site and search field size) we found neither tDCS-specific (anodal > sham) visual search improvements, nor stimulation-specific baseline dependencies (larger improvements for lower baseline performers were observed in both tDCS conditions, suggesting rather stimulation-unspecific effects like regression to the mean). Together, our results provide evidence against reliable top-down attention-guided visual search improvements through FEF tDCS.

## Main

Top-down attention, the voluntary (de-)prioritization of information like the selection of certain visual sub-fields [1] controlled by a dorsal fronto-parietal network including the frontal eye field (FEF) [2], is central to successful everyday-life behavior and poses immense problems when negatively impacted by disease [3,4]. This relevance makes it a key target for stimulation-based enhancement and rehabilitation attempts in healthy and clinical populations, respectively [5,6]. Transcranial direct current stimulation (tDCS) has been proposed as safe and easy tool for non-invasively manipulating neural excitability (and thereby possibly cognition) through the scalp-level application of weak electric currents [7]. In line with this, a recent study reported significant visual search improvements through anodal (excitatory) vs. sham (control) tDCS over the left FEF (lFEF) [8]. This effect was inversely related to baseline performance (larger stimulation-specific improvements for low baseline performers) [8]. Based on the proposed dominance of the right FEF (rFEF) for visual-spatial top-down attention [9], in a first experiment, we tested whether the previously reported visual search improvement [8] could be reproduced and potentially further augmented through rFEF instead of lFEF stimulation.

All experiments were preregistered and will provide open data/analyses upon publication. For experimental details please refer to this article’s supplement. In experiment one, *N*=29 healthy volunteers (*M*_*age*_=21.62, *SD*_*age*_=1.82; female=15, male=12, diverse=2) performed a visual search task, in which they indicated the presence or absence of an upside-down ‘T’ amongst upright ‘T’s and ‘L’s, before (baseline) or during (peri) the application of sham or anodal rFEF tDCS. Only in the anodal but not the sham condition, the rFEF was stimulated throughout the task using constant 2 mA currents with the anode placed over rFEF and the cathode placed above the left eye.

To test our hypothesis of larger stimulation-related visual search improvements through anodal vs. sham rFEF tDCS, we computed a one-tailed paired-sample *t*-test with stimulation (anodal/sham) as independent variable and baseline-corrected percentage correct as dependent variable. The results of experiment one are depicted in row one of Figure 1. Although there was good overall task performance without ceiling effect (sham-baseline: *M*=78.41, *SD*=6.74; sham-peri: *M*=78.24, *SD*=7.74; anodal-baseline: *M*=78.69, *SD*=5.19; anodal-peri: *M*=80.00, *SD*=6.33), the preregistered *t-*test was far from being significant (sham: Δ*M*=-0.17, Δ*SD*=7.76; anodal: Δ*M*=1.31, Δ*SD*=6.18; *t*(28)=0.67, *p*=.253, *d*=0.125). Exploratory Bayesian statistics demonstrated about three times more evidence for *H*_*0*_ over *H*_*1*_ (*BF*_*10*_^*+*^=0.36). Moreover, while the previous finding of an inverse relationship between baseline performance and tDCS-related changes [8] could be reproduced in exploratory correlation analyses of the anodal condition (ρ(27)=-0.44, *p*_*Bonf*_=.017), the same was true for the sham condition (ρ(27)=-0.51, *p*_*Bonf*_=.005).

**Fig. 1.**
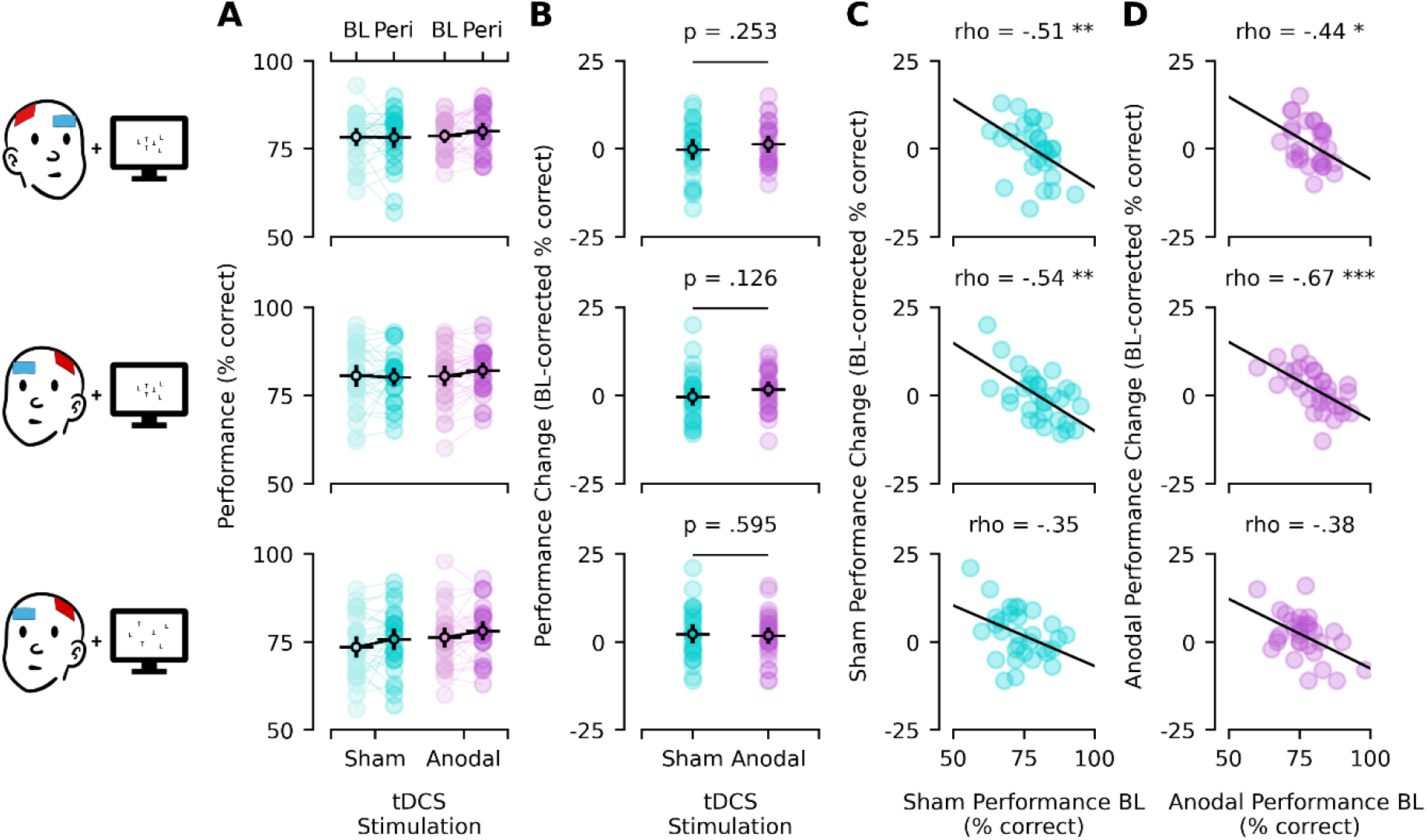
**(A)** Visual search performance before and during sham or anodal FEF tDCS, **(B)** stimulation-related performance changes relative to baseline and **(C**,**D)** their association with baseline performance separately for sham and anodal tDCS, respectively. Rows relate to the different experiments/conditions (row one: experiment one, rFEF tDCS, small search fields; row two: experiment two, lFEF tDCS, small search fields; row three: experiment two, lFEF tDCS, large search fields). Colored dots represent single-subject data, whereas black circles with horizontal bars represent sample means with 95% confidence intervals (vertical bars). *Note: Electrode setups and search fields were simplified for illustration purposes; p-/rho-values refer to the respective t/Spearman test results with asterisks indicating statistical significance, where* * ≙*p* < .05, ** ≙*p* < .01 and *** ≙*p* < .001; *BL = baseline*.

Our first experiment could, thus, neither boost nor reproduce the previously reported visual search improvement following lFEF tDCS [8] and suggests that apparent stimulation-related effects (improvements for low baseline performers and vice versa) might reflect rather stimulation-unspecific effects (regression to the mean). However, one could argue that (in contrast to our hypothesis and previous literature) rFEF tDCS might have been less suitable for this task than the previously used lFEF stimulation [8]. Moreover, compared to the original study [8], we used rather small search fields (7.2° visual angle vs. whole-screen presentations), which might require smaller spatial attention shifts and shorter saccades, thereby potentially limiting visual search improvements through FEF stimulation. Therefore, in a second experiment, we applied lFEF tDCS (mirroring electrode positions from experiment one) before and while *N*=31 new participants (*M*_*age*_=21.48, *SD*_*age*_=2.31; female=22, male=9, diverse=0) searched either small (7.2° x 7.2°) or large (21.3° x 13.1°) visual fields (the procedure itself remained unchanged).

To test our hypothesis of search field size as the most probable potential limitation in experiment one (and, hence, to test our expectation of greater visual search improvements through anodal vs. sham lFEF tDCS for large but not small search fields), we repeated the above *t-*test separately for small and large visual fields. The results of experiment two are depicted in rows two (small fields) and three (large fields) of Figure 1. In line with experiment one, performance was overall good without ceiling effects, but again largely comparable across tDCS sessions and test times for both search field sizes (sham-baseline_small_: *M*=80.55, *SD*=8.50; sham-peri_small_: *M*=80.16, *SD*=7.01; anodal-baseline_small_: *M*=80.45, *SD*=7.96; anodal-peri_small_: *M*=82.10, *SD*=6.41; sham-baseline_large_: *M*=73.52, *SD*=8.27; sham-peri_large_: *M*=75.77, *SD*=8.53; anodal-baseline_large_: *M*=76.29, *SD*=7.80; anodal-peri_large_: *M*=78.10, *SD*=7.30). Again, the preregistered *t-*tests were far from being significant (sham_small_: Δ*M*=-0.39, Δ*SD*=7.00; anodal_small_: Δ*M*=1.65, Δ*SD*=5.82; *t*(30)=1.17, *p*=.126, *d*=0.210; sham_large_: Δ*M*=2.26, Δ*SD*=7.17; anodal_large_: Δ*M*=1.81, Δ*SD*=6.35; *t*(30)=-0.24, *p*=.595, *d*=-0.044). Exploratory Bayesian statistics provided 2-6 times more evidence for *H*_*0*_ over *H*_*1*_ (small: *BF*_*10*_^*+*^=0.62; large: *BF*_*10*_^*+*^=0.16). For small search fields the negative correlation between baseline performance and tDCS-related changes was once more reproduced but again for both anodal and sham tDCS (anodal: ρ(29)=-0.67, *p*_*Bonf*_<.001; sham: ρ(29)=-0.54, *p*_*Bonf*_=.003), whereas for large search fields exploratory correlation analyses failed to reach significance in any condition (anodal: ρ(29)=-0.38, *p*_*Bonf*_=.069; sham: ρ(29)=-0.35, *p*_*Bonf*_=.108).

Our second experiment thus supports our previous null-effect despite lFEF stimulation and large search fields [8] and renders alternative explanations for such null-effect (specifically rFEF stimulation and small search fields) rather unlikely. Together, our two experiments therefore provide evidence against reliable stimulation-specific visual search improvements through FEF tDCS. It should be noted, however, that we cannot exclude potential effects of rFEF stimulation in combination with large search fields, nor can we entirely exclude other experimental differences as potential explanations for diverging results (no formal replication). Even in the case of a true null-effect, this conclusion might be limited to this specific task and/or to healthy young populations with well-tuned attention systems more generally. One potential obstacle to consistent tDCS-related visual search improvements might lie in the method’s low spatial resolution: Single-cell FEF stimulation in monkeys has shown improved detection for stimuli within the cell’s receptive field but impairments for information outside of it [10], suggesting complex and potentially conflicting effects through widespread tDCS stimulation. Thus, whereas the present experiments do not support substantial visual search improvements through tDCS in healthy young participants, FEF stimulation might prove fruitful for attentional improvements with spatially more fine-tuned methods and/or in clinical/elderly populations with suboptimal attention tuning and greater potential for cognitive enhancement.

## CRediT authorship contribution statement

**CP:** Conceptualization, Data curation, Formal analysis, Investigation, Methodology, Project administration, Resources, Software, Supervision, Validation, Visualization, Writing – original draft, Writing – review and editing; **CA, RGH, JR:** Investigation, Methodology, Resources, Writing – review and editing; **MAD, ME, MPG, FL, SM, CR:** Investigation, Methodology, Writing – review and editing; **JT:** Investigation, Resources, Writing – review and editing; **PS:** Conceptualization, Formal analysis, Funding acquisition, Investigation, Methodology, Project administration, Resources, Software, Supervision, Validation, Writing – review and editing

## Acknowledgements

We thank the LMU students (Anastasia Bauer, Larissa Behnke, Benedikt Biroga, Vico Bittorf, Christina Dietz, Niko Erdösi, Elena Habelt, Ruth Henneke, Annika Hofmann, Nia Nedialkova, Veit Peteranderl, Rebekka Rau, Julius Russ, Franziska Salzberger, Carolin Straub, Eirini Thomopoulou, Liv Tilgner, Christopher Williams, Anastasia Witt, Lea Zucker) for their support with study planning and/or execution.

