## Supplement for "Transcranial direct current stimulation over the frontal eye field has no effect on visual search performance"

### Participants

A preregistered *a-priori* power analysis in G*Power© (v3.1.9.4; Heinrich Heine University, Düsseldorf, Germany; Faul et al., 2007) for a one-tailed paired-sample *t*-test with an estimated effect size of Cohen's *d*=0.5 (as conservative approximation of effect sizes reported by previous studies with FEF tDCS effects on different visual search outcomes; Diana et al., 2022; Gan et al., 2022), an alpha level of α=0.05 and a desired power of 1-β=0.80 indicated a required sample size of *N*=27. Based on this estimate and a potential drop-out of participants over the course of our two-session experiments, we tested *N*=29 and *N*=32 participants in experiments one and two, respectively. In contrast to our preregistered policy of no performance-based participant exclusions, we excluded *n*=1 participant from experiment two *post-hoc* due to unreasonably low scores across all conditions of the sham session (all <22% correct), suggesting a potential confusion of the stimulus-response mapping. The final sample of experiment two therefore consisted of *n*=31 participants.

All participants were right-handed according to the Edinburgh Handedness Inventory (Oldfield, 1971), reported normal or corrected-to-normal vision and no history of neurological or psychological disorders and fulfilled all tDCS safety inclusion criteria (no diagnosis or close family history of epilepsy, traumatic brain injuries, concussions/blackouts, tinnitus, pregnancy, metallic implants or cochlear/neural/cardiac stimulators, excitability-increasing medication, spinal cord operations or cerebrospinal puncture, sleep deprivation or drug/increased alcohol intake within 24 hours before the experiment). Participants gave their written informed consent at experiments' beginnings (privacy rights were met throughout) and could receive course credits as compensation at their ends. The studies were approved by LMU's F11 ethics committee (ethics approval reference number '02_2022_Sauseng_a' from April 6th 2022) and performed according to laws and the declaration of Helsinki.

### Visual Search Task

The visual search task was presented on a standard laptop using Presentation® (v0.7; Neurobehavioral Systems©, Berkeley, CA, USA) and consisted of a visual search field (1750 ms; 3000 ± 500 ms inter-trial interval), comprising a total of 30 letters (0.4° visual angle height each): 15 upright 'L's, 14 upright 'T's and either an upside-down 'T' (target present trial, 50%) or an additional upright 'T' (target absent trial, 50%). The participants' task was to freely explore the visual search field (i.e., without restriction of eye movements) and to indicate target presence or absence via button press as quickly and precisely as possible ('Y'/'N' using the index finger of the left/right hand, respectively).

The visual search task in experiments one and two only differed with respect to search field size and trial count/task duration: In experiment one, the search field was always small (7.2° x 7.2° visual angle) and each assessment (baseline/peri) consisted of one single block with 60 trials (30 target absent/present trials; conditions randomly interleaved) and lasted about 4.75 min, adding up to a total duration of about 35 min per session (sham/anodal) including training (20 trials) and tDCS preparation. In experiment two, only half of the trials contained a small search field (7.2° x 7.2° visual angle), whereas the other half of trials contained a large search field (21.3° x 13.1°). To account for the additional field size condition while maintaining comparability with the first experiment, here, each assessment (baseline/peri) consisted of one single block with 120 trials (30 target present/absent in small/large search field trials; conditions randomly interleaved) and lasted about 9.5 min, adding up to a total duration of about 50 min per session (sham/anodal) including training (40 trials) and tDCS preparation.

### tDCS Stimulation

In both experiments, stimulation via constant current was delivered using a TCT Research Limited (Hong Kong, China) tDCS device. Positioning of the two saline-soaked sponge electrodes (5 cm x 5 cm each) closely resembled that of Gan and colleagues (2022) with the anode placed over the FEF (4 cm anterior and 5 cm right/left lateral to the vertex for rFEF/lFEF stimulation in the first/second experiment, respectively) and the cathode placed above the opposite eye. For the anodal stimulation, current was ramped up to 2 mA (20 sec), maintained at that level for the duration of the visual search task and then ramped back down to 0 mA (20 sec). For the sham stimulation, current ramped up to 2 mA (20 sec) and then immediately ramped back down to 0 mA (20 sec) (similar to Gan et al., 2022). To allow potential stimulation-driven excitability changes to unfold before task execution (Gan et al., 2022), participants received five minutes of passive stimulation before starting the visual search task for the peri assessment. The two types of stimulation (sham/anodal) were delivered in two separate single-blinded sessions at least 24 hours apart with session order counterbalanced across participants.

### Stimulation Blinding

To assess blinding success (and hence participant expectations and potential response biases), at the end of both experiments' second session, participants were asked to assign stimulation types to test sessions (in experiments one and two) and to indicate expected stimulation effects (excitation/inhibition; in experiment two only). Although most participants correctly assigned stimulation types to test sessions (experiment one: 28/29 participants, 96.55%; experiment two: 26/31 participants, 83.87%), only 14/31 (45.16%) participants from experiment two correctly identified the intended stimulation effect.

### Analysis

Performance in each condition (sham/anodal, baseline/peri, small/large search fields) was computed in Excel (Microsoft®, Redmond, Washington, USA) and later analyzed in Jamovi (v2.3.28; Jamovi Developer Team, 2024) as baseline-corrected percentage correct (percentage correct during stimulation minus percentage correct before stimulation, where each trial with a matching first response was considered correct). To account for non-normality and/or outliers, which were present in some of the variables used for the correlation but not the group-level mean difference analyses, we computed parametric *t*-tests (incl. Bayesian equivalents) and non-parametric Spearman rank correlations. Bonferroni correction was used to account for multiple correlation testing. Results were visualized in PowerPoint (Microsoft®, Redmond, Washington, USA) and the Spyder environment (v5.4.3; Spyder Developer Team, 2020) for Python (v3.11.3; Van Rossum & Drake, 2009) using custom-written scripts and the open-source packages NumPy (v1.24.3; Harris et al., 2020) and Matplotlib (v3.7.1; Hunter, 2007).
